# Neuronal loss of *Galnt2* in Mice Impairs O-glycosylation and Recapitulates Neurological Features of GALNT2-CDG

**DOI:** 10.1101/2024.09.30.615951

**Authors:** Andrew C. Edmondson, Melody Yu, Alvin Villarosa, Emily J. Shiplett, John Hintze, Sergey Y. Vakhrushev, Katrine T. Schjoldager, Zhaolan Zhou

## Abstract

GALNT2-CDG is a multi-system genetic disorder due to biallelic pathogenic mutations in *GALNT2*, which encodes a ubiquitously expressed, Golgi-localized glycosyltransferase that initiates mucin-type O-glycosylation. Affected individuals exhibit dysmorphic facial features, short stature, decreased HDL-C, and notable impairments in brain function. GALNT2-CDG patients show global developmental delay without speech development, childhood epilepsy, autistic-like features, and white-matter brain abnormalities. However, the role of O-glycosylation in brain development and function remains poorly understood. To address this question, we selectively ablated *Galnt2* from neuronal cells in the brain and found that conditional knockout mice exhibit deficits across numerous behavioral domains, including locomotion, motor coordination, sociability, learning, and memory, as well as spontaneous seizures, recapitulating characteristic neurological manifestations of GALNT2-CDG. Given the catalytic activity of GALNT2 in initiating mucin-type O-glycosylation, we used glycoproteomics to identify alterations in O-glycosylation in synaptosomes purified from cortical tissues. We ascertained a non-redundant, isoenzyme-specific contribution of GALNT2 to the cortical synaptosomal O-glycoproteome, revealing candidate glycoproteins and disrupted O-glycosites associated with behavioral abnormalities in knockout mice. Together, these findings demonstrate a functional role of O-glycosylation in neurons, implicating O-glycosylation in diverse molecular and cellular pathways important for neuronal function and provide new insights into the neurological pathophysiology of GALNT2-CDG.

## Introduction

Congenital disorders of glycosylation (CDG) are a group of multi-system genetic disorders that disrupt cellular glycosylation machinery. Most CDG identified to date disrupt N-glycosylation, however, some additionally or exclusively disrupt O-glycosylation (3). Patients can exhibit symptomatic dysfunction of essentially any organ, but most also exhibit neurological deficits of varying severity (4). These disorders emphasize the importance of glycosylation as an essential posttranslational modification, yet the the mechanisms underlying neurological dysfunction in CDG remain poorly understood.

Mucin-type or *N*-acetyl-D-galactosamine (GalNAc)-type O-glycosylation (subsequently O-glycosylation) is a common post-translational modification predicted to occur on more than 80% of proteins traversing the secretory pathway (5). O-glycosylation is initiated by a large family of polypeptide GalNAc-transferase (GALNT, also known as GalNAc-T) isoenzymes with substantial overlapping protein substrate specificities. Aberrant function of a single isoenzyme from the GALNT gene family generally results in loss of a minor subset of non-redundant protein O-glycosylation (6). Nevertheless, these changes can have substantial health impacts, such as in the case of GALNT3, where autosomal recessive loss causes familial tumoral calcinosis (7).

*GALNT2* encodes the ubiquitously expressed Golgi-localized GALNT2 isoenzyme and was initially implicated in human lipoprotein metabolism (8, 9). The cause of the lipid abnormalities was shown to be due to non-redundant GALNT2-mediated O-glycosylation of angiopoietin-like 3 (ANGPTL3), apolipoprotein C-III (apoC-III) (10), and phospholipid transfer protein (PLTP) (1). GALNT2-CDG presents as a multi-system genetic disorder due to biallelic pathogenic mutations in *GALNT2* (11). Affected individuals exhibit dysmorphic facial features, short stature, and decreased High Density Lipoprotein Cholesterol (HDL-C), however, the most impactful disease symptoms are neurological. GALNT2-CDG patients exhibit global developmental delay without speech development, childhood epilepsy, autistic-like features, and white-matter brain abnormalities on magnetic resonance imaging (11). These symptoms suggest that O-glycosylation is critical to neurodevelopmental functions, and while recent progress has expanded our knowledge of the neuronal O-glycoproteome (12, 13) the specific contribution of individual GALNT isoenzymes and impact on the pathophysiology remains unknown.

The pathogenic variants resulting in GALNT2-CDG abolish germline GALNT2 enzymatic activity, and initial studies correlating rodent whole body *Galnt2* knock-out (KO) model phenotypes with GALNT2-CDG patient phenotypes demonstrated recapitulation of multisystemic phenotypic features, including poor growth, low HDL-C, and behavioral/neurodevelopmental abnormalities (11). These behavioral phenotypes included increased exploratory behavior in open field activity assessment and impairments of motor coordination on accelerating rotarod and sociability on social choice procedure (11). However, this model exhibits substantial embryonic lethality (less than half the expected numbers of KO mice are born (11)) and reduced growth of surviving pups raising concerns regarding the representational validity of studying only surviving mice of this model and systemic confounds for behavioral testing.

To address these questions, here we bred a floxed *Galnt2* allele (*Galnt2^fl/fl^*) (1) with a post-mitotic pan-neuronally expressing Cre driver mouse line, Snap25-IRES2-cre (*Snap25^Cre/+^*) (14, 15) to produce *Galnt2* neuronal knock-out (nKO) mice. Through behavioral assays and electroencephalogram (EEG) recordings of brain activity we demonstrate patient relevant neurological deficits, including motor, sensory, learning and memory functions as well as spontaneous seizures in nKO compared to floxed littermate controls. We hypothesized that observed abnormalities could be localized to the neuronal synapse and lead to impaired synaptic transmission. Therefore, we used O-glycoproteomics to ascertain the non-redundant, isoenzyme-specific contribution of GALNT2 to the cortical synaptosomal O-glycoproteome and map site-specific protein glycosylation, providing a basis for investigating how O-glycosylation modulates synaptic structure and function.

## Results

### Generation and molecular validation of conditional Galnt2 nKO mice

The conditional *Galnt2^fl^* mouse allele (1) contains LoxP sites surrounding exons 4-6. These exons contribute to the catalytic domain of GALNT2 and their cre-mediated excision results in frameshift with premature termination of the coding transcript and nonsense-mediated decay (Figure 1A). We used a polymerase chain reaction (PCR) based genomic recombination assay to confirm Cre-mediated recombination of LoxP sites and excision of exons 4-6 in the cortex of nKO mice (Figure 1B). We also used quantitative PCR (qPCR) to quantify *Galnt2* expression in nKO mice in cortex and cerebellum (Figure 1, C and D, respectively), confirming significantly decreased expression of *Galnt2* in the brain of nKO mice. Residual expression is likely due to RNA from non-Cre expressing cells, such as glia and other non-neuronal cells in the brain. Unfortunately, we were unable to acquire a reliable antibody to quantify the reduction in GALNT2 protein in the brain via Western blot.

**Figure 1.**
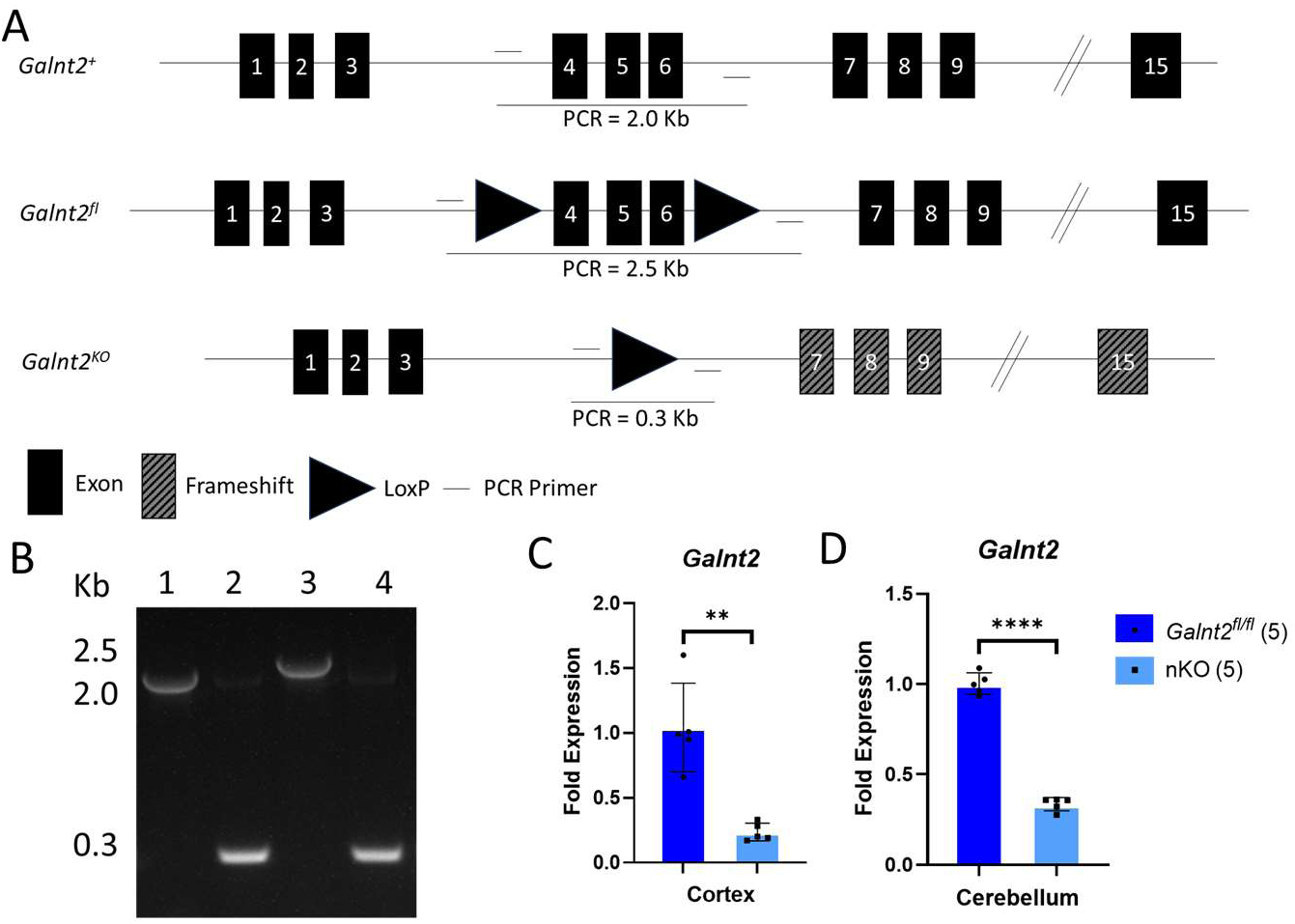
Molecular characterization of *Galnt2* nKO mice. (A) Schematic of mouse genomic loci of *Galnt2^+^* (WT), *Galnt2^fl^* (conditional KO) and *Galnt2^KO^* (KO) alleles with PCR strategy indicated. (B) Genomic Recombination Assay PCR of cortical DNA from 1, WT; 2, KO; 3, Conditional KO; 4, nKO. (C and D) qPCR quantitation of *Galnt2* expression in cortex and cerebellum, respectively. Number of biological replicates indicated in parentheses and analyzed using Welch’s t-test. **, *P* < 0.01; ****, *P* < 0.0001 Alt text: Data on *Galnt2* DNA and RNA in mouse model brain tissue with subfigures labelled A to D, illustrating genomic recombination as detected by PCR and statistical analyses of gene expression.

### Characterization of Galnt2 nKO mice

nKO mice were bred by crossing heterozygous floxed *Galnt2* and heterozygous *Snap25-IRES2-*Cre females (*Galnt2^fl/+^; Snap25^Cre/+^*) with homozygous floxed *Galnt2* males (*Galnt2^fl/fl^; Snap25^+/+^*) to avoid rare cre-mediated germline recombination of the *Galnt2* floxed allele in male mice (data not shown) and generate cohorts of mice for further evaluation. Resulting genotypes of 150 pups across 27 litters showed expected genotype ratios of ∼25% nKO mice (*P* = 0.3, χ^2^ *test*), suggesting no embryonic lethality. Some of the mice studied were generated by breeding homozygous floxed *Galnt2* and heterozygous *Snap25-IRES2-*Cre females (*Galnt2^flfl^; Snap25^Cre/+^*) with homozygous floxed *Galnt2* males (*Galnt2^fl/fl^; Snap25^+/+^*) and achieving ∼50% nKO mice (data not shown).

Mice were monitored and weighed weekly. Both male and female nKO mice demonstrated excessive weight gain with aging compared to littermate controls, with *P* < 0.001 for genotype and *P* < 1.0 x 10^-15^ for week x genotype interaction for females and *P* < 0.05 for genotype and *P* < 0.05 for week x genotype interaction for males using two-way matched values ANOVA (Figure 2A and B, respectively). Adult brain weight appeared slightly larger in female nKO mice compared to littermate controls (*P* < 0.05), with similar trend identified in male nKO mice (*P* = 0.055) (Figure 2 C and D, respectively). Given abnormal body weight and weight correlations with metabolic syndrome including lipid abnormalities, and the association of GALNT2 with lipid abnormalities (8, 9), fasting lipids were assessed in adult male mice between 16 and 18 weeks of age. No lipid abnormalities were identified (Supplemental Figure 1), consistent with GALNT2’s role in lipid metabolism primarily being restricted to the liver (1), rather than also having a prominent central nervous system (CNS) contribution. A complete clinical veterinary pathological assessment via necropsy was completed on 7 floxed littermate females, 7 nKO females, 4 floxed littermate males, and 6 nKO males between 18-20 weeks of age. Necropsy confirmed excess weight in nKO female mice compared to littermate controls with similar trend in male mice (Supplemental Figure 2A). Necropsy also suggested slightly larger liver and kidneys in female nKO mice compared to littermate controls, without similar trend identified in male nKO mice (Supplemental Figure 2B-F). No other macroscopic, microscopic, or histopathologic abnormalities correlating with genotype were identified in the liver, kidneys, brain or adipose tissue (Supplemental Figure 3A-H), suggesting phenotypic abnormalities in nKO mice arise from a functional rather than structural etiology.

**Figure 2.**
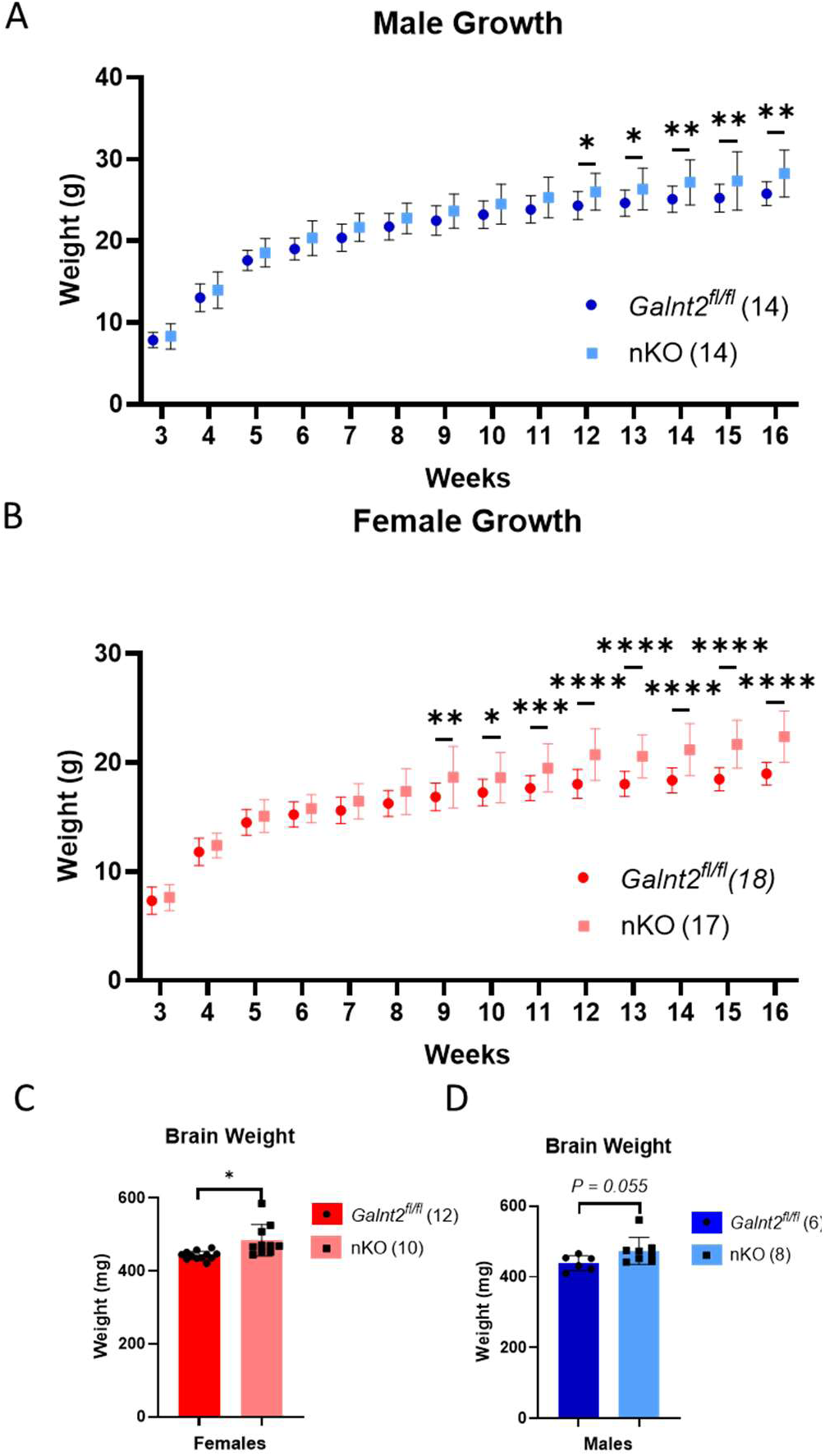
Body and brain weights of *Galnt2* nKO mice. Weekly body weights of (A) male and (B) female mice grouped by genotype. Growth charts initially analyzed using two-way matched values ANOVA. Pairwise comparisons at each timepoint analyzed using Fisher’s Least Significant Differences test. Brain weights of (C) female and (D) male mice, grouped by genotype. Analyzed using Welch’s t-test. Values plotted as mean values, error bars represent SD. Number of biological replicates indicated in parentheses. *, *P* < 0.05; **, *P* < 0.01; ****, *P* < 0.0001. Alt text: Data on mouse model body and brain weights with subfigures labelled A to D, illustrating increased body and brain weights in nKO mice.

### Neuronal loss of Galnt2 induces broad neurobehavioral deficits

To systematically characterize the contribution of neuronal GALNT2 to GALNT2-CDG relevant behavioral phenotypes we performed a battery of behavioral tests (Figure 3A) over a 2-month period starting at 10-12 weeks of age and found broad deficits across neurological motor, sensory, learning, and memory domains in both male and female adult mice. Consistent with observations of GALNT2-CDG patients, males and females are similarly affected, although with some sex-specific differences noted on behavioral testing. Results for combined analysis are presented in Figure 3 and results of sex-separated analysis is presented in Supplementary Figures 4 and 5.

**Figure 3.**
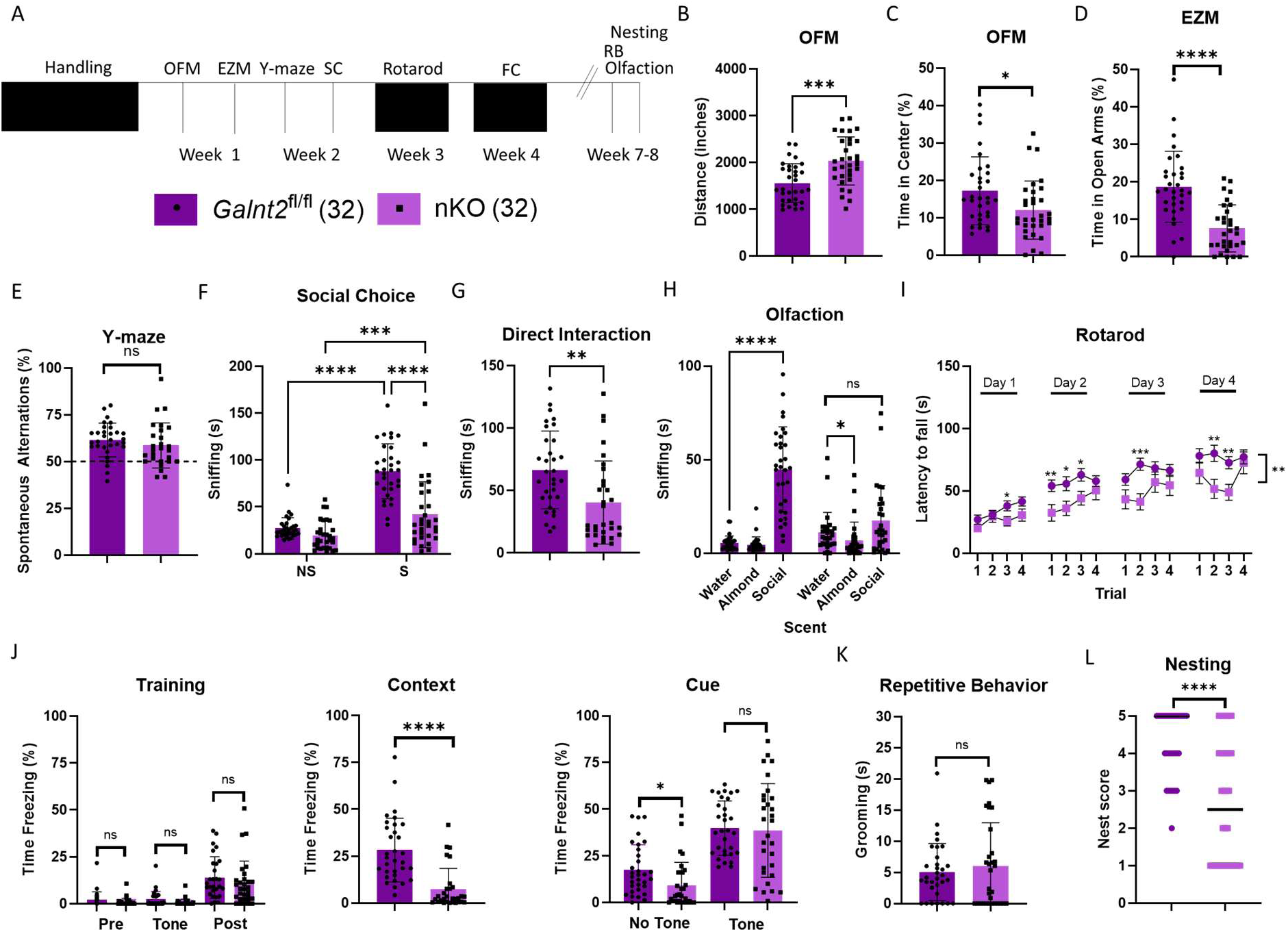
*Galnt2* nKO mice broad deficits across neurological motor, sensory, learning, and memory domains. Timeline of behavioral testing battery illustrated in (A) with legend. (B) nKO mice explore the OFM more than floxed littermates, but (C) spend less time in the center of the OFM. (D) nKO spend less time in the open arms of the EZM. (E) nKO have intact short-term and spatial working memory in the Y-maze. (F) nKO spend less time than floxed littermates sniffing and (G) directly interacting with a novel stimulus mouse during the 3-chambered SC test, but (H) this could also be due to lack of social scent discrimination. (I) nKO fall more quickly from an accelerating, rotating rod than floxed littermates. (J) nKO mice spend significantly less time freezing compared with floxed littermates when returned to the fear-conditioning testing chamber (contextual) but (I) the same amount of time freezing upon hearing the footshock-associated tone (cue). (K) nKO mice spend no more time in repetitive grooming than floxed littermates. (L) nKO have trouble building well formed nests. For all panels, floxed, *n* = 32; nKO, *n* = 32. Welch’s unpaired, 2-tailed t-test, except as follows: 3-chambered SC assay and olfaction, 2-way ANOVA with Fisher’s Least Significant Differences test; rotarod, generalized linear mixed effects model with Geisser-Greenhouse correction and Fisher’s Least Significant Differences test; nesting, unpaired Mann-Whitney test. *, *P* < 0.05, **, *P* < 0.01, ***, *P* < 0.001, ****, *P* < 0.0001. Data are represented as mean ± SD, except rotarod where errors bars represent ± Standard Error of the Mean (SEM) and nesting where the line indicates median. Alt Text: Data from mouse behavioral analysis with subfigures labelled A to L, illustrating statistical analyses of deficits across neurological motor, sensory, learning, and memory domains in *Galnt2* nKO mice.

Given developmental delays among GALNT2-CDG patients (11), we first assessed nKO motor locomotor function and exploratory behavior in the open field maze (OFM) (16) where mice demonstrated increased distance traveled and decreased time spent in the center of the maze (Figure 3B and 3C), results typical of an anxiety-like phenotype. The anxiety-like phenotype was further confirmed with the elevated zero maze (EZM) where nKO mice spent less time in the open arms of the maze (Figure 3D). Short-term and spatial working memory were assessed with the Y-maze (17) with notable difference between sexes, while not significantly impaired in the combined analysis (Figure 3E), male nKO mice had significant deficits (Supplemental Figure 4D) while female nKO mice did not (Supplemental Figure 5D). Given the autistic-like behaviors of GALNT2-CDG patients, we assessed social behavior in nKO mice using the three-chambered social choice (SC) test (18), where each animal explores cylinders containing either a novel, inanimate object (NS, non-social cue) or a novel conspecific mouse (S, social cue), followed by a period of time where the mouse is allowed to directly interact with the novel conspecific mouse. Consistent with social deficits, nKO mice interacted less with the conspecific mouse during both the SC (Figure 3F) and direct interaction (Figure 3G) phases of the test.

However, this must be interpreted with caution as they also demonstrated a potential deficit in social scent discrimination, not spending significantly more time sniffing social scent than water (Figure 3H). On accelerating rotarod testing (19), nKO mice showed deficits in coordination with reduced latency to fall (*P* < 0.01 for genotype factor, Figure 3I). We next assessed learning and memory in the fear conditioning paradigm (20) and identified a deficit in context-specific memory deficit with intact cue-specific memory (Figure 3J). Another characteristic autistic-like behavior is repetitive behaviors, which were assessed in nKO mice as excessive grooming and found to be normal (Figure 3K). Finally, in an assessment of complex task integration, nKO mice demonstrated a deficit in being able to construct nests (Figure 3L).

### In Vivo EEG reveals seizure activity and cognitive impairment in Galnt2 nKO mice

While handling mice for behavioral assays, occasionally abnormal movements in some mice were noted, including rapid head jerks, tonic stiffening, and forelimb clonus, which were suggestive of behavioral seizures. Additionally, given that seizures affect GALNT2-CDG patients, we performed surgical intracranial electrode placement and recorded brain electrical activity at the cortical surface and within the brain in the region of the hippocampus via *in vivo* EEG with simultaneous video recording for an average of approximately 78 hours/mouse (range 42.6 – 113.4 hours). We recorded *in vivo* EEG in a cohort of 8 younger nKO mice (mean age ∼12 weeks, range 7.9 – 16 weeks) and a cohort of 8 older nKO mice (mean age ∼35 weeks, range 30 – 44 weeks), as well as 7 older wild-type (WT) age-matched littermate controls. Seizures captured during the recording period were typically stage 3 (clonic seizure with forelimb clonus and rearing) or stage 4 (tonic-clonic seizure with loss of righting reflex), according to the modified Racine scale (21, 22). An EEG tracing from a typical seizure event is depicted in Figure 4A. We observed an apparent increase in seizure prevalence in the older cohort of mice with 3/8 (38%) younger nKO mice and 7/8 (88%) of older nKO mice demonstrating seizures during the recording period.

**Figure 4.**
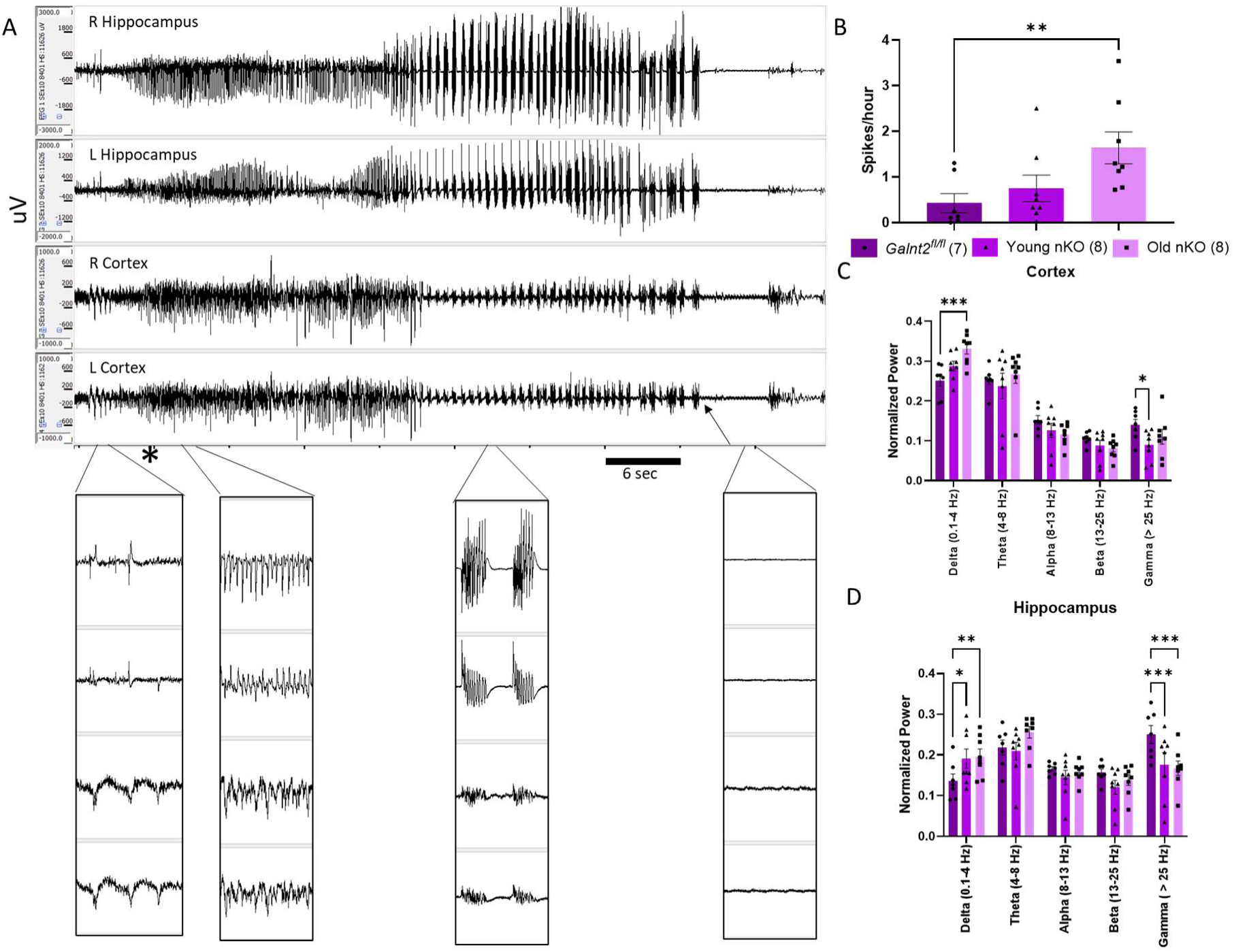
*Galnt2* nKO mice demonstrate seizures and altered background electrical activity on in vivo EEG. (A) 1 minute of EEG tracing demonstrating a typical *Galnt2* nKO seizure with 1 second insets at indicated phases of the seizure demonstrating spiking, spike-wave, polyspike wave, and suppression patterns, respectively. * indicates behavioral start of seizure, arrow indicates burst-suppression. *Galnt2* nKO mice demonstrate (B) increased epileptiform spikes as well as altered EEG background frequency band composition in the (C) cortex and (D) hippocampus. Results from the entire recording period for each mouse were averaged for each animal shown as an individual data point. Results were analyzed by one-way ANOVA for spikes and two-way ANOVA by band and genotype, followed by Fisher’s Least Significant Differences test to compare between groups. *, *P* < 0.05, **, *P* < 0.01, ***, *P* < 0.001, ****, *P* < 0.0001. Data are shown as mean ± SEM. Alt Text: Data from mouse *in vivo* EEG analysis with subfigures labelled A to F, illustrating EEG tracings and statistical analyses of EEG background spiking and frequency band composition in *Galnt2* nKO mice.

All mice demonstrating seizures had at least 2 seizures identified during their recording period. None of the 7 floxed littermate control mice were identified to have a seizure. The average amount of recording time of mice identified to have seizures compared to mice not having seizures during the recording period was not significantly different (younger mice with seizures, 95 hours; younger mice without seizures, 72 hours; older mice with seizures, 72.4 hours; older mice without seizures, 87 hours).

We assessed for electrographic spikes and altered EEG background frequency composition in nKO mice compared with floxed control mice using power analysis. All mice were evaluated without regard to whether they had an observed seizure during the recording period, however, any seizure events were excluded from EEG background analysis. *Galnt2* nKO mice demonstrate increased spike burden (Figure 4B) and a shift in background activity toward faster frequencies (Figure 4, C and D).

Specifically, nKO mice demonstrate decreased relative power in the slow delta frequency band (0.1-4 Hz) and increased relative power in the faster gamma frequency band (> 25 Hz). Despite the increased seizure prevalence in older nKO mice, the EEG background frequency composition was consistent in both the younger and older cohorts of nKO mice, although spike burden was only significantly increased in the older nKO mice.

### Ascertaining the GALNT2-specific Cortical Synaptosomal O-glycoproteome

Based on the altered neuronal network dynamics identified on *in vivo* EEG in nKO mice and the seizure phenotype of GALNT2-CDG patients, we hypothesized that loss of neuronal GALNT2 could affect glycoproteins in the neuronal synapse that might contribute to the development or propagation of seizures. We isolated enriched cortical synaptosomes from the brains of nKO and floxed littermate mice (Supplemental Figure 6) as we previously described for WT mice (12). By using multiplex tandem mass-tag (TMT) amine-reactive labelling (23) of tryptic peptide digests from synaptosomal preparations followed by Jacalin lectin weak-affinity chromatography (LWAC), we generated comparative O-glycoproteomes from nKO and floxed littermates (Figure 5A) and determined the non-redundant, isoenzyme-specific contribution of GALNT2 to the cortical synaptosomal O-glycoproteome. Although male and female GALNT2-CDG patients and mice are similarly affected, we performed comparative O-glycoproteomics in male and female mice separately, using the separate experiments to demonstrate reproducibility and increase confidence in our findings. We performed liquid chromatography-tandem mass spectrometry (LC-MS/MS) proteome analysis of the LWAC flow-through where we identified 2,412 TMT-labelled peptides across 759 proteins in female and 2,584 TMT-labelled peptides across 659 proteins in male synaptosomes. We did not identify any significantly altered protein abundances across the synaptosomal proteomes from *Galnt2* nKO compared to floxed controls (Supplemental Figure 7).

**Figure 5.**
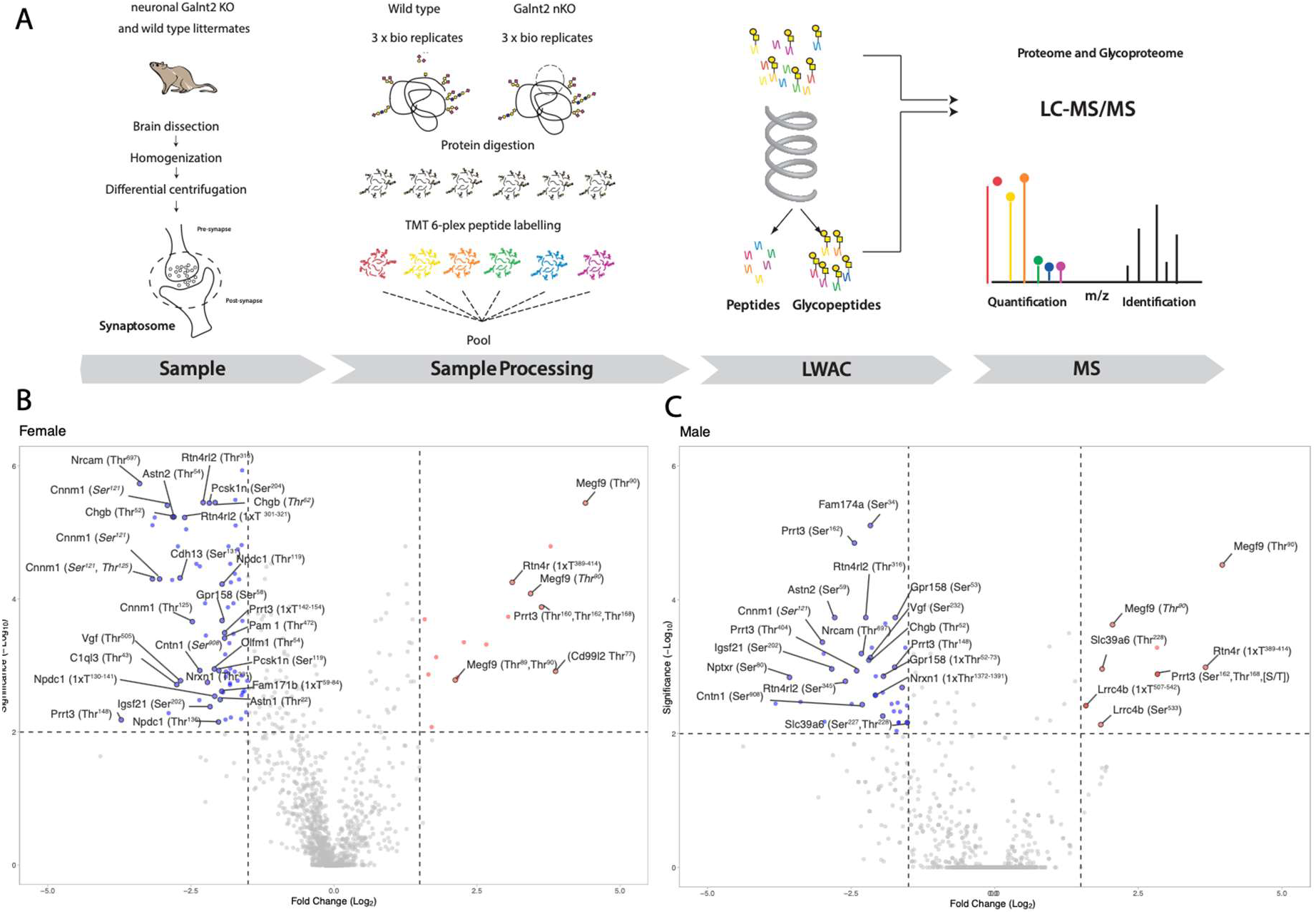
Non-redundant, isoenzyme-specific contribution of GALNT2 to the cortical synaptosomal O-glycoproteome. (A) Schematic of differential O-glycoproteomic workflow in synaptosomes. Differential glycoproteome between *Galnt2* nKO and floxed littermates is plotted as volcano plots of nKO/control fold change for (B) female and (C) male synaptosomes. Proteins are pseudocolored and labelled for fold change > 3 and *P* < 0.01, blue, decreased glycosylation in nKO; red, increased glycosylation in nKO. Alt Text: Data from O-glycoproteomic evaluation of synaptosomes with subfigures labelled A to G, illustrating results.

The jacalin-lectin enriched glycopeptides were analysed by LC-MS/MS to produce GALNT2 differential O-glycoproteomes (Table 1, Figure 5, B and C). We identified 1,172 TMT-labelled O-glycopeptides across 190 glycoproteins from the female synaptosome preparations and 1,298 TMT-labelled O-glycopeptides across 205 glycoproteins from the male synaptosome preparations. There was a 68% overlap of identified proteins and 77% overlap of identified glycoproteins between female and male synaptosome preparations. At a nominal significance level of *P* < 0.05, we identified 290 O-glycopeptides from 84 glycoproteins with differential abundance between *Galnt2* nKO mice and floxed littermates in female synaptosomes (Figure 5B) and 122 differential O-glycopeptides from 39 glycoproteins in male synaptosomes (Figure 5C). The majority of differentially abundant O-glycopeptides represent loss of site-specific glycosylation in *Galnt2* nKO mice, as would be anticipated for a non-redundant, isoenzyme-specific contribution of GALNT2. At sites with gain of O-glycosylation, this likely represents an alternative glycosylation performed by another GALNT isoenzyme when GALNT2-mediated O-glycosylation is not present or the increased abundance of a glycopeptide containing multiple glycosites where one site is GALNT2 specific. Comparing differential O-glycopeptides between the male and female cohorts, there were 13 glycopeptides from 9 glycoproteins with replication of the exact glycopeptide in the same direction in both cohorts, representing highly confidence non-redundant, isoenzyme-specific contributions of GALNT2 to the cortical synaptosomal O-glycoproteome (Table 1).

**Table 1.**
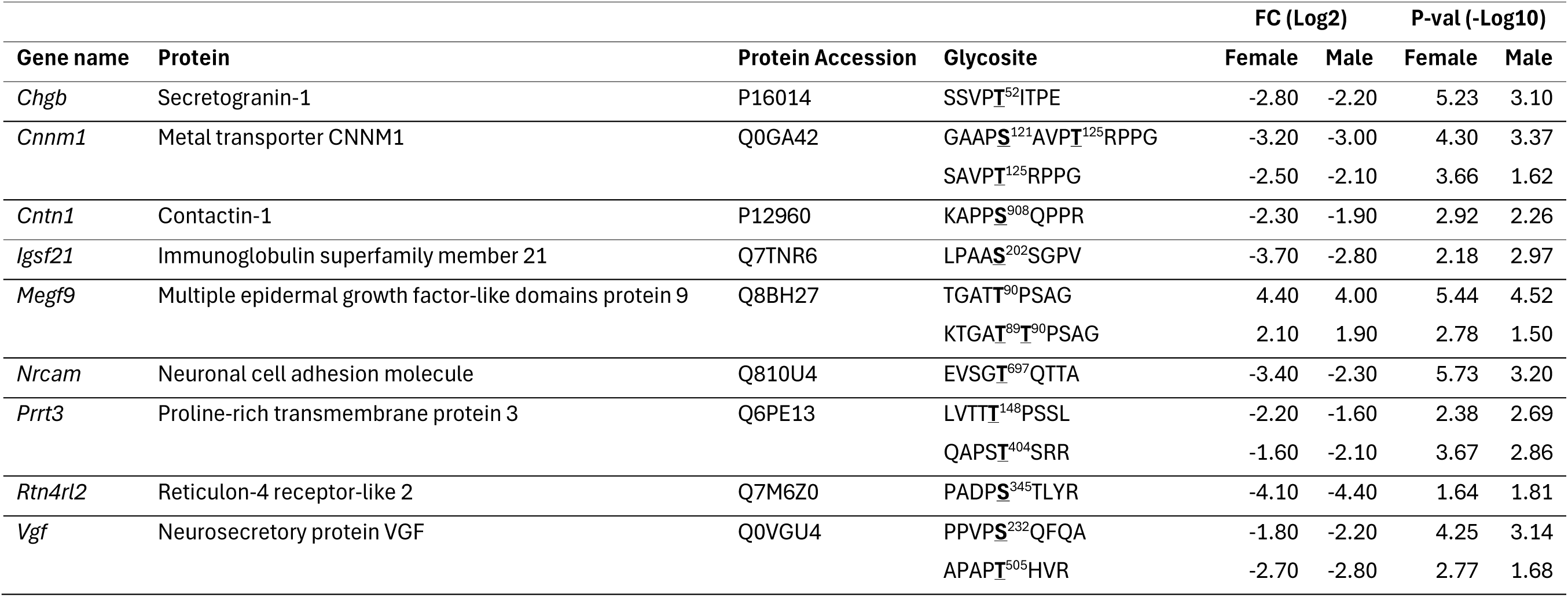
Non-redundant, isoenzyme-specific contribution of GALNT2 to the cortical synaptosomal O-glycoproteome.

## Discussion

GALNT2-CDG is a genetic disorder with multi-system manifestations, including severe involvement of the nervous system resulting in global developmental delay without speech development, epilepsy, and autistic-like features (11). We previously attempted to utilize a whole-body KO mouse model to better understand the disorder (1, 11), however, substantial embryonic lethality and the multisystem impacts of GALNT2 loss raised concerns about the representational validity of surviving mice and the ability to interpret neurologic involvement given systemic confounds. Here we validated a post-mitotic pan-neuronal conditional *Galnt2* KO mouse model to rigorously interrogate and understand the neurological features of GALNT2-CDG.

We demonstrate that mice without GALNT2 in neurons exhibit patient-relevant neurological deficits across motor, sensory, learning, and memory domains. The nKO mice also recapitulate a key neurological deficit of GALNT2-CDG, spontaneous seizures. GALNT2-CDG patients have been described with a range of seizure types and severities, including febrile seizures, infantile spasms, and multifocal epilepsy with frequent treatment resistance (11). Outside of the seizures recorded in our *Galnt2* nKO mice, interictal EEG analysis demonstrated frequent abnormal electrographic spikes and background differences in EEG frequency composition. Similar abnormalities have been observed in many epilepsy models. The background EEG frequency composition changes in *Galnt2* nKO mice, specifically the shift toward high-frequency background activity, is an EEG abnormality associated with cognitive impairment (24).

Our model does have some important limitations. Outside of direct impacts due to GALNT2 deficiency, seizures could also impact neurodevelopment and thus our behavioral assessments. Individual mouse behavioral assessments were excluded if there was a concern for seizure activity during the assessment. Our behavioral characterization exhibited some discrepancies from what was previously observed in whole body KO mice, including differences in anxiety-like phenotype, motor function on rotarod testing, and olfaction (11). It is unclear to what extent the results were impacted by representational validity or experimental differences in behavioral testing. Additionally, our nKO model retains GALNT2 in glia and therefore fails to elucidate the role of GALNT2 in glial cells, such as astrocytes. It is also possible that intact GALNT2 in glia may modify the neurological assessments, explaining some of the differences we observe in *Galnt2* nKO mice from the whole body KO mice.

Further, despite our efforts to avoid systemic confounds present in *Galnt2* whole body KO mice (11), the *Galnt2* nKO mice exhibited increased weight gain that could influence their performance on some motor behavioral tasks. Interestingly, GALNT2 was recently implicated in a brain-liver neurocircuit where knock-down of *Galnt2* specifically in the ventromedial hypothalamus resulted in significantly increased body weight, food intake, and blood glucose levels compared with controls (25), positioning GALNT2 in the central control of metabolic homeostasis, which is mirrored in our nKO mouse phenotype. Future efforts to characterize non-redundant contributions of GALNT2 to the O-glycoproteome in the ventromedial hypothalamus will provide further valuable insights into glycoprotein participants in central metabolic homeostasis. A discrepancy between nKO or ventromedial hypothalamic knock-down of *Galnt2* and whole body KO, where the mice are smaller than WT littermates (11), suggests GALNT2 also plays additional roles in growth in the periphery. This observation is further corroborated by recent reports of GALNT2’s role in pancreas development and function using conditional transgenic overexpression of GALNT2 (26).

We also demonstrated that the neuronal synapse, an area enriched in glycoproteins, relies on GALNT2 for glycosylation of key synaptic proteins. We identified many non-redundant, isoenzyme-specific contributions of GALNT2 to the cortical synaptosomal O-glycoproteome. The 13 glycosites we identified with high confidence were replicated across two experiments. However, there are likely additional GALNT2-reliant glycosites that were not replicated or entirely missed due to technical limitations. As our model also retains GALNT2 in astrocytes, protein contributions from astrocytes in our synaptosome isolations could also have masked some GALNT2-reliant glycosites.

Our findings implicate a role for O-glycosylation in diverse neurological processes, including learning, memory, and neurotransmission. Consistent with most other cells and tissues we have analyzed to date (1, 27), GALNT2 displays a minor non-redundant, isoenzyme-specific contribution to the O-glycoproteome in synaptosomes without altering protein abundances. Thus, the subset of target glycoproteins we identified displaying differential O-glycosylation are key candidates for explaining the observed neurological deficits in our nKO mice. Glycoproteomic analysis of isolated synaptosomes suggest GALNT2 supports synapse formation or stability (via cell adhesion molecules such as CNTN1, NRCAM, IgSF21, and RTN4FL2) and synaptic transmission (such as mediated by secretory granules with CHGB or neuropeptides including VGF) (Figure 6). Additional studies defining the functional consequences of these glycosylation events on protein functions will provide important insights into pathophysiology of neurological deficits in GALNT2-CDG, as well as the roles of these glycosylation events in normal neuronal function.

**Figure 6.**
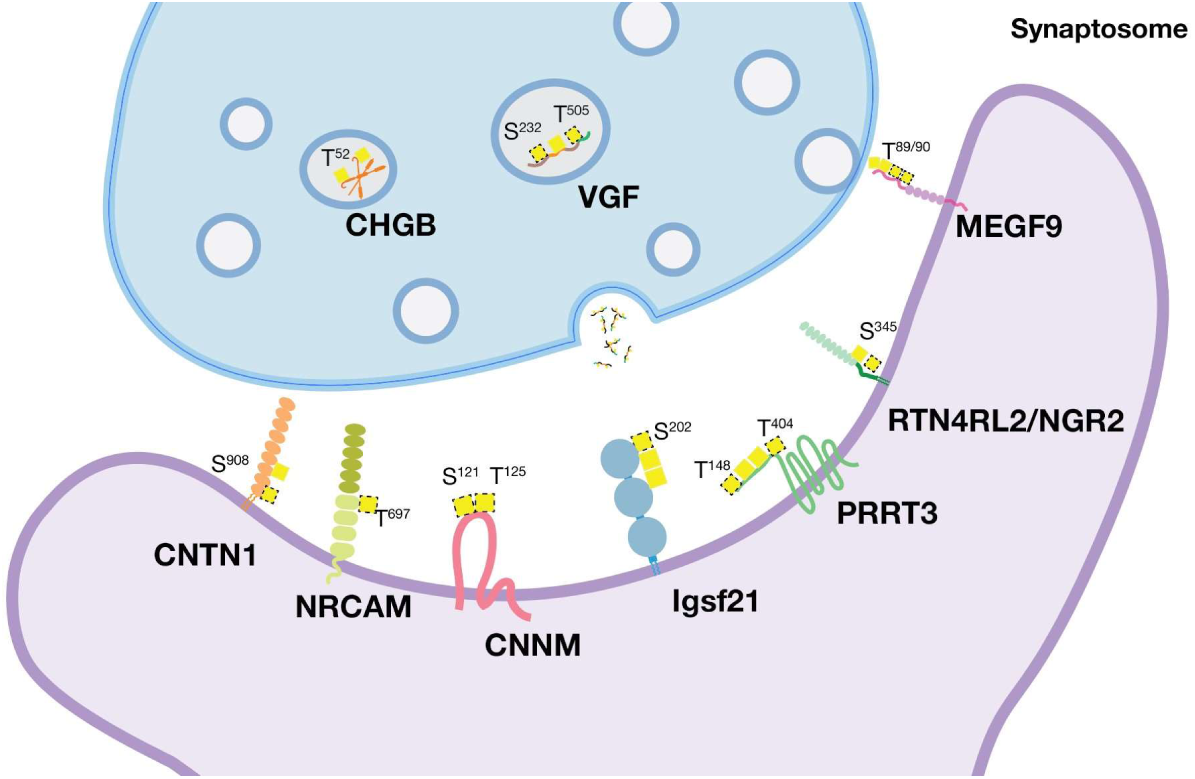
Non-redundant, isoenzyme-specific contribution of GALNT2 in the cortical synapse. Depiction of the cortical synapse with focus on GALNT2-initiated O-glycosylation. GALNT2-initiated glycosylation indicated by dashed GalNAc (yellow square) outline. Monosaccharide symbols follow the Symbol Nomenclature for Glycans system (2). Alt Text: Depiction of the cortical synapse showing synaptic proteins with GALNT2-initiated O-glycosylation from Table 1.

## Methods

### Sex as a biological variable

Our study examined male and female animals, and similar findings are reported for both sexes with sex-dimorphic effects reported when identified.

### Mouse strains and genotyping

The *Galnt2* conditional KO line was previously described (1) and are available from Taconic Biosciences (*Galnt2* model 10264). Snap25-IRES2-Cre is available from the Jackson Laboratory (strain no. 023525). *Galnt2* conditional KO mice were a kind gift from Dr. Daniel J. Rader at the University of Pennsylvania. Snap25-IRES2-Cre mice were obtained from Jackson Laboratory. All lines were maintained in the C57Bl/6J background after being back-crossed more than 10 generations. All progeny were genotyped either with the previously reported PCR strategies (1) or with qPCR by Transnetyx, Inc (Cordova, TN).

### Animal Husbandry

Mice were group housed in cages of 2 to 5 on a 12-hour light/12-hour dark cycle with food and water provided ad libitum. Each breeding cage contained one male (*Galnt2^fl/fl^; Snap25^+/+^*) and one to two females (*Galnt2^fl/+^; Snap25^cre/+^* or *Galnt2^fl/fl^; Snap25^cre/+^*) Male littermates and female littermates were weaned at 3 weeks of age and housed in sex-segregated, mixed genotype cages.

### Molecular validation

Genomic DNA and RNA was isolated from mouse cortex using the AllPrep DNA/RNA purification kit (Qiagen, Germantown, MD). Genomic recombination assay was performed using PCR primers that flank exons 4-6 of mouse Galnt2 (F: 5’-GTACGTGAGACAGGCCTAAGG-3’ R: 5’-CAAGCTTCATTTAGGACCAAGC-3’) and EmeraldAmp PCR Master Mix (Takara Bio, San Jose, CA). PCR cycling parameters were as follows: 1 cycle of 95°C for 30 seconds; 35 cycles of 95°C for 30 seconds, 60°C for 30 seconds, 72°C for 2.5 min; and 1 cycle of 72°C for 10 min. cDNA was made from isolated RNA using the High Capacity cDNA Reverse Transcription Kit (Applied Biosystems, Foster City, CA) according to the manufacturer’s instructions. TaqMan gene expression assays (Applied Biosystems, Foster City, CA) were used to detect mouse *Galnt2* exon 6 to 7 junction (Mm00519805_m1) and *Hprt* (Mm03024075_m1). Reactions contained 0.5 µL 20x assay mix, 5 µL 2x TaqMan reaction mix, and 1 µL cDNA and were performed in triplicate. Quantitative RT-PCR was performed on an Applied Biosystems 7900HT real-time PCR system. Relative expression differences were calculated by the delta delta Ct method using *Hprt* as the housekeeping gene.

### Systemic Evaluation

Mice were weighed weekly. Serum from adult mice for lipid analysis was collected via retro-orbital bleed from mice fasted from food for 4 hours with continued access to water. Lipids were measured by IDEXX Bioanalytics (North Grafton, MA). Complete clinical veterinary necropsy was performed by University of Pennsylvania Comparative Pathology Core collecting organ weights, formalin-fixing and paraffin embedding organs, sectioning, and staining with hematoxylin and eosin following standard clinical procedures. The pathological assessment was performed by trained veterinary pathologists in a blinded fashion without knowledge of the experimental group distribution and genotype of the animals.

### Behavioral analysis

Mouse behavioral testing was performed in mice beginning at 10-12 weeks of age. Mice were habituated to handling for two weeks prior to starting behavioral testing. Mice were habituated to the testing room for at least 1 hour before testing (except for context-and cue-dependent fear-conditioning assays), which was always performed at the same time of day. Testing was performed in the same order for each animal, beginning with OFM assay, then the EZM assay, Y-maze, 3-chambered SC assay, accelerating rotarod assay, context-and cue-dependent fear conditioning, followed by two weeks recovery, then repetitive behavior assay, overnight nesting, and olfaction, as diagramed in Figure 3A. See Supplemental Methods for details on all behavioral assessments. Investigators were blinded to genotype while conducting and scoring behavioral assays. Mice that exhibited behavior concerning for a seizure (e.g. rapid head jerks, tonic stiffening, forelimb clonus) during an assay or that failed to engage in the assay (e.g. did not move from their initial placement in the testing apparatus) were excluded from analysis for that assay.

### Four-Channel EEG implantation surgery

Mice were stereotaxically implanted with an electrode assembly under continuous isoflurane anesthesia. The electrode assembly consisted of 6 leads: 2 surface electrodes attached to miniature skull screws placed over the left and right frontal cortices (from Bregma: A/P +1.0 mm, M/L ± 1.5 mm); 2 depth coated electrodes in left and right hippocampus (from Bregma: A/P −2.2 mm, M/L ± 2.0 mm, D/V +1.7 mm); and a reference and a ground electrode also attached to miniature skull screws directly behind Lambda on either side of midline (from Bregma: A/P – −5.2 mm, M/L ± 1.3 mm);. Silver wires (0.13 mm diameter) were attached to each electrode and connected to an 8-pin headmount (Pinnacle Technology Inc, Lawrence, KS). The entire assembly was secured on the skull with dental cement (Ortho-Jet, Lang Dental, Wheeling, IL). Mice were given at least 72 hours post-surgery to recover prior to being placed in a Plexiglas recording cage (Pinnacle Technology Inc, Lawrence, KS). EEG waveform was amplified by a preamplifier with a gain of 10 pV (Pinnacle Technology Inc, Lawrence, KS) at the head of the animal before being passed through a low-torque commutator to the Sirenia Acquisition system (Pinnacle Technology Inc, Lawrence, KS) for final-stage amplification and filtering. EEG was sampled at 2000 Hz in a 12-hour light/dark cycle with food and water provided ad libitum. Video was watched by a reviewer trained to identify seizures. EEG traces were processed with the application of a 1 Hz high-pass filter, 500 Hz low-pass filter, and 60 Hz notch filter and visualized within Serenia Seizure (Pinnacle Technology Inc, Lawrence, KS). To confirm the quality of our EEG recordings, we administered 100 mg/kg pentylenetetrazol (PTZ) subcutaneously at the conclusion of video monitoring. All mice exhibited typical PTZ-induced seizures including sharp spikes, spike-wave, and polyspike discharges, typical characteristic of PTZ-induced seizures.

### EEG Analysis

EEG was analyzed using MATLAB with code adapted from (28) and is available at <u>github.com/avrosa-md/GALNT2-CDG-EEG-Code</u>. Analysis steps included: 1) Detection of poor-quality recording channels: An automated artifact detector was used on each recording channel before analysis. Detection of a noisy or poorly recording channel was completed by calculating the average of the root mean squared amplitude and skew of the voltage for each second of the recording. Channels with a root mean square amplitude of less than 30 μV or more than 200 μV or a skew greater than 0.4 were excluded from further analysis. 2) Spike detection: A spike detector algorithm designed to detect voltage deflections that were greater than 5 standard deviations above the mean with a full width at half maximum amplitude of 5–200 ms was used to detect spikes. Spikes were eliminated if they occurred within 10-seconds of a deflection found to be an artifact (i.e., it had a half width outside the 5-to 200-ms range). Multi-channel spikes were defined as a spike occurring in at least 2 channels. 3) Power analysis: To analyze alterations in background EEG frequency composition in *Galnt2* nKO mice, recordings were divided into 5-second epochs for power analysis. Epochs containing large amplitude artifacts, most likely due to movement (defined as a peak Z-score of the root mean squared voltage amplitude greater than 3), were excluded from frequency analysis. Fast Fourier transform analysis was then completed on artifact-free epochs. Quantification of the power in each of the major EEG frequency bands (delta 0.1–4.0 Hz, theta 4–8 Hz, alpha 8–13 Hz, beta 13–25 Hz, and gamma 25–50 Hz) was averaged across artifact-free epochs within 10-minute recording windows. Power in each band across all epochs was averaged and power in each band was normalized to total power.

### Synaptosome isolation

We isolated synaptosomes from mouse cortex using modifications of previously published protocols for Percoll gradient isolation (29) and synaptosome protein preparation for mass spectrometry analysis (30). Briefly, we homogenized ∼200 mg of flash frozen cortical tissue in 2 ml of isotonic sucrose solution (0.32 M Sucrose, 1 mM EDTA, 5 mM Tris-HCl, pH 7.4, 0.25 mM DTT) containing protease inhibitors (Roche, Little Falls, NJ) using a Dounce glass homogenizer, centrifuged at 1000g for 10 min at 4℃ to remove nuclei. The supernatant was diluted to 2 ml with isotonic sucrose solution and applied to a discontinuous Percoll gradient (3% vol/vol, 10% vol/vol, 15% vol/vol, and 23% vol/vol). The Percoll gradient was then centrifuged at 31,000g for 5 min at 4℃. Fractions 3 and 4 were combined, diluted to 50 ml with isotonic sucrose solution, centrifuged at 20,000g for 30 min at 4℃ to remove Percoll. The resulting pellet was resuspended in acetone and the proteins precipitated via incubation in ice-cold acetone at -20℃ overnight. Proteins were isolated by centrifugation at 15,000g for 30 min at 4℃, acetone removed, and pellet allowed to air dry. Protein pellets were stored at -80 until proteomic and glycoproteomic analysis.

### O-Glycoproteomics

The synaptosomal O-glycoproteome was recently reported (12), however only data from WT mice was presented. Here we present a quantitative differential O-glycoproteome analysis of TMT-labelled glycopeptides from *Galnt2* nKO compared to WT littermates.

### nLC-MS/MS analysis

LC–MS/MS was performed on a system composed of an EASY-nLC 1000 (Thermo Fisher Scientific) interfaced via a nanoSpray Flex ion source to an Orbitrap Fusion/Lumos Tribrid (Thermo Fisher Scientific), equipped for both higher energy collisional dissociation (HCD) and electron transfer dissociation (ETD) modes, enabling peptide sequence analysis with and without retention of glycan site-specific fragments, respectively as previously described (12). For the quantitative analysis, only HCD MS2 spectra were used. In the case of ETD MS2 spectra, the group of TMT reporter fragment ions (*m*/*z* range of 126–132) was extracted from the adjacent HCD MS2 spectrum (the same precursor ions), concatenated with the corresponding ETD MS2 spectrum, and later used for quantification. Processing of the TMT MS3 data was performed using Proteome Discoverer version 2.1 software (Thermo Scientific).

### Visualization of MS data

Lists of quantified O-glycopeptides from TMT-labelled glycopeptides from WT and *Galnt2* nKO synaptosome preps are presented in Table 1. O-glycopeptides with more than one missing quant values were dropped. Data was normalized to synaptosome O-glycopeptides from WT mice and the ratios were log_2_-transformed. The data is presented in a volcano plot displaying the log_2_ of the fold change on the x-axis and minus log_10_ of the p-value on the y-axis.

### Statistics

For statistical analyses we used Welch’s unpaired, 2-tailed t-test, except as follows: genotype distributions, χ2 test against expected Mendelian distribution; growth charts and rotarod, two-way matched values ANOVA with pairwise comparisons at each timepoint analyzed using Fisher’s Least Significant Differences test; 3-chambered SC assay and olfaction, two-way ANOVA with Fisher’s Least Significant Differences test; nesting, unpaired Mann-Whitney test; spikes, 1-way ANOVA; EEG background frequency band composition, two-way ANOVA with Fisher’s Least Significant Differences test. All graphs are plotted using Prism (GraphPad). Level of significance was set at *P* ≤ 0.05. In our figures, * is used to denote all 0.01 < *P* < 0.05, ** for 0.001 < *P* < 0.01, *** for 0.0001 < *P* < 0.001, and **** for *P* < 0.0001. Data are represented as mean ± SD, except rotarod and EEG analysis where errors bars represent ± SEM and nesting where the line indicates median.

## Study Approval

Experiments were conducted in accordance with the ethical guidelines of the National Institutes of Health and with the approval of the Institutional Animal Care and Use Committee of the University of Pennsylvania.

## Data Availability

*Data underlying this article will be shared on reasonable request to the corresponding author*.

## Author Contributions

ACE, KTS, and ZZ designed research studies. ACE, MY, AV, and KTS performed statistical analysis. ACE, JH, and SYV conducted experiments. ACE, EJS, JH, and SYV acquired data. ACE, MY, AV, and KTS analyzed data. ACE wrote the manuscript. ACE, MY, AV, EJS, JH, SYV, KTS, and ZZ revised the manuscript.

## AI disclosure

The authors declare that no AI tools were used in the original research for or writing of this article.

## Abbreviations

ANGPTL3: angiopoietin-like 3
apoC-III: apolipoprotein C-III
CDG: Congenital Disorders of Glycosylation
CNS: central nervous system
EEG: electroencephalogram
ETD: electron transfer dissociation
EZM: Elevated Zero Maze
FC: fold change
GalNAc: *N*-acetyl-D-galactosamine
GALNT: GalNAc-transferase
HDL-C: High Density Lipoprotein Cholesterol
HCD: higher energy collisional dissociation
KO: knock-out
LWAC: lectin weak-affinity chromatography
LC-MS/MS: liquid chromatography-tandem mass spectrometry
nKO: neuronal knockout
ns: non-significant
NS: non-social
OFM: Open Field Maze
PTZ: pentylenetetrazol
PCR: polymerase chain reaction
PLTP: phospholipid transfer protein
qPCR: quantitative polymerase chain reaction
RB: repetitive behavior
S: Social
SC: Social Choice
SD: Standard Deviation
SEM: Standard Error of the Mean
TMT: tandem mass-tag
WT: wild-type

