## Supplemental materials for "Neuronal loss of *Galnt2* in Mice Impairs O-glycosylation and Recapitulates Neurological Features of GALNT2-CDG"

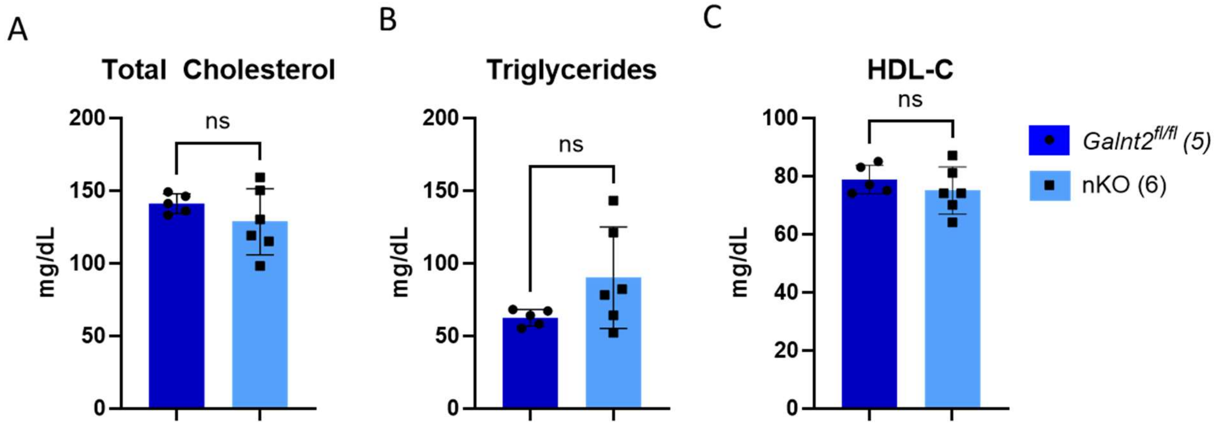


**Supplemental Figure 1. Fasting blood lipids in *Galnt2* nKO mice.** Four-hour fasting measurements of A) total cholesterol, B) triglycerides, and C) HDL-C. LDL-C was also measured and below the detectable limits of the assay in all mice. Analyzed using Welch's t-test. Values plotted as mean values, error bars represent SD. Number of biological replicates indicated in parentheses. ns, non-significant.

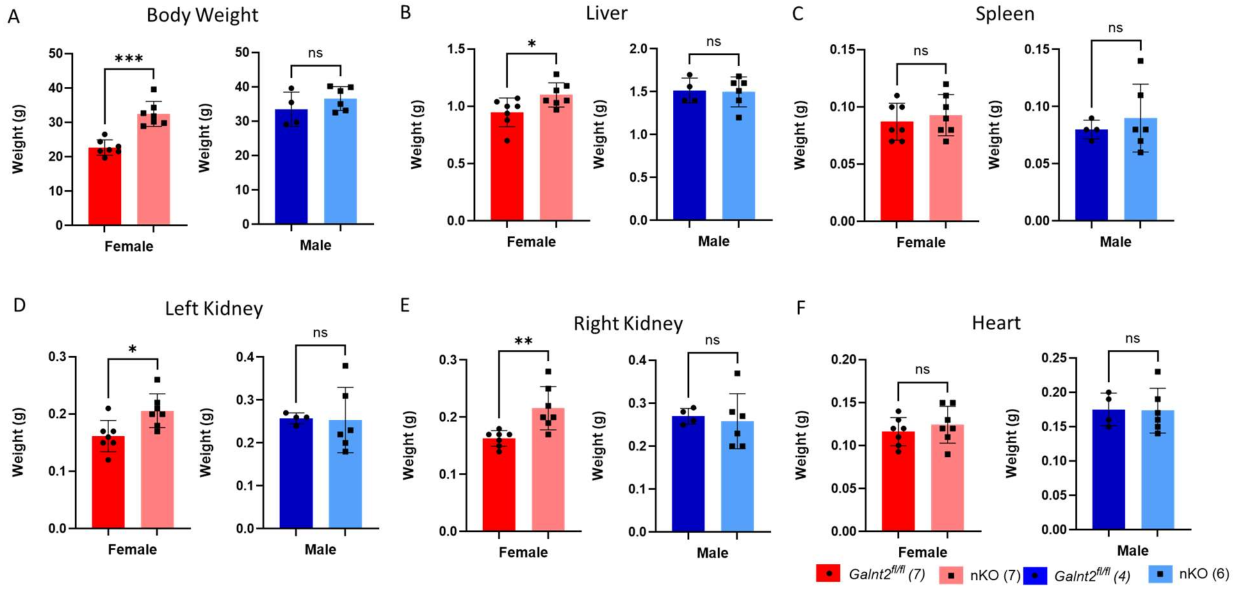


**Supplemental Figure 2. Necropsy weights of *Galnt2* nKO mice.** Weights of female and male mice, respectfully, grouped by genotype for A) body weight, B) Liver, C) Spleen, D) Left Kidney, E) Right Kidney, and F) Heart. Analyzed using Welch's t-test. Values plotted as mean values, error bars represent SD. Number of biological replicates indicated in parentheses. ns, non-significant, *, $P < 0.05$; **, $P < 0.01$; ***, $P < 0.001$.

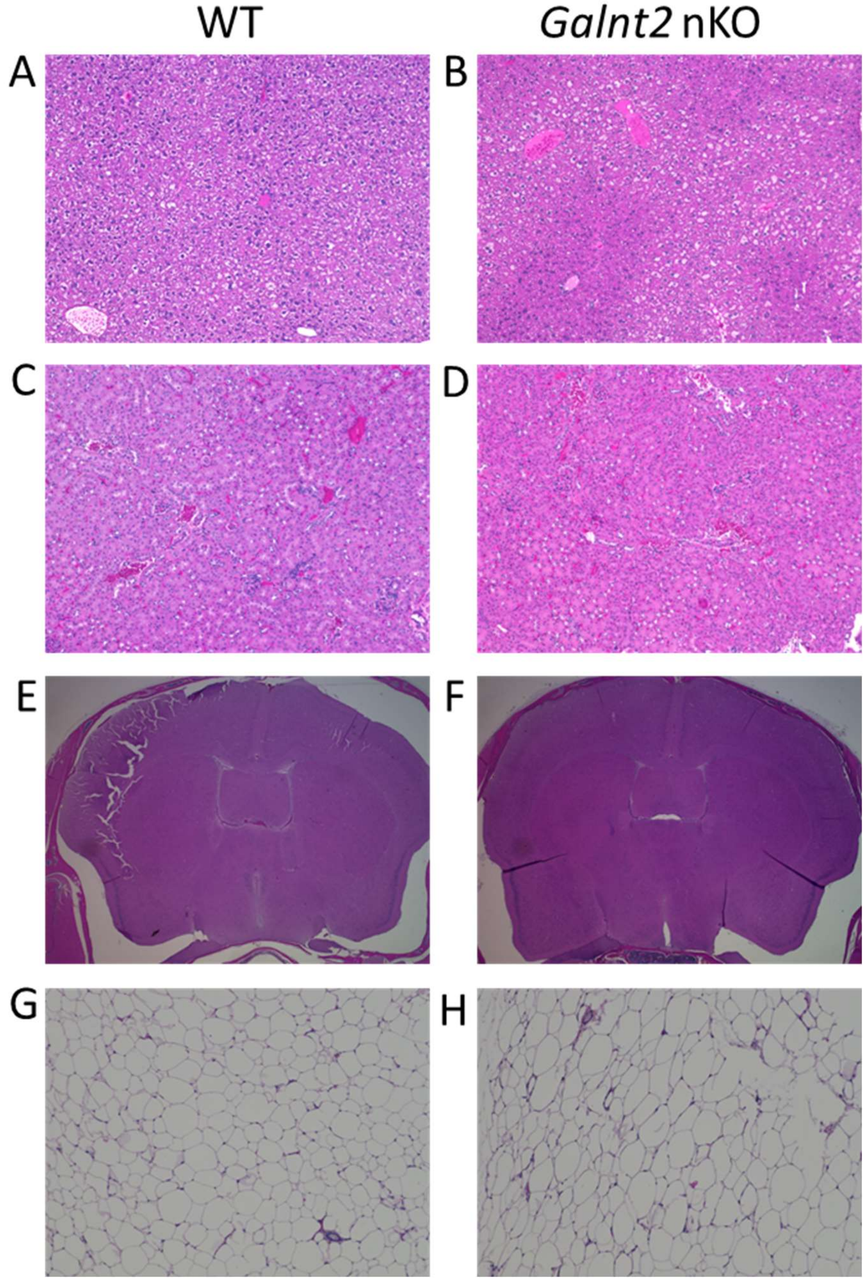


**Supplemental Figure 3. Organ histology of *Galnt2* nKO mice.** Representative images of H&E stained liver (A,B), kidney (C,D), brain (E,F), and adipose (G,H) from WT and *Galnt2* nKO mice, respectfully.

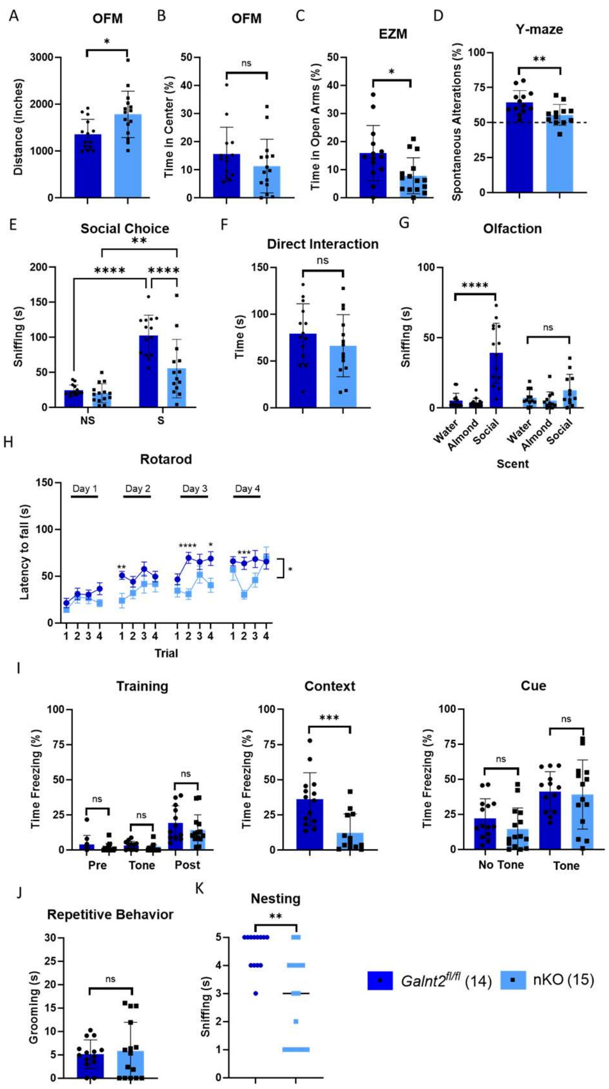


**Supplemental Figure 4. Male *Galnt2* nKO mice exhibit broad deficits across neurological motor, sensory, learning, and memory domains.** (A) Male nKO mice explore the OFM more than floxed littermates with (B) similar time spent in the center of the OFM. (C) Male nKO spend less time in the open arms of the EZM. (D) Male nKO have impaired short-term and spatial working memory in the Y-maze. (E) Male nKO mice spend less time than floxed littermates sniffing but similar amount of time (F) directly interacting with a novel stimulus mouse during the 3-chambered SC test, which could be due to (G) lack of social scent discrimination. (H) Male nKO fall more quickly from an accelerating, rotating rod and take longer to learn the task than floxed littermates. (I) Male nKO mice spend significantly less time freezing compared with floxed littermates when returned to the fear-conditioning testing chamber (contextual) but (I) the same amount of time freezing upon hearing the footshock-associated tone (cue). (J) Male nKO mice spend no more time in repetitive grooming than floxed littermates. (K) Male nKO mice have trouble building well-formed nests. For all panels, floxed, $n$ = 14; nKO, $n$ = 15. Welch's unpaired, 2-tailed t-test, except as follows: 3-chambered SC assay and olfaction, two-way ANOVA with Fisher's Least Significant Differences test; rotarod, two-way matched values ANOVA with Fisher's Least Significant Differences test; nesting, unpaired Mann-Whitney test. *, $P < 0.05$, **, $P < 0.01$, ***, $P < 0.001$, ****, $P < 0.0001$. Data are represented as mean ± SD, except rotarod where errors bars represent ± SEM and nesting where the line indicates median.

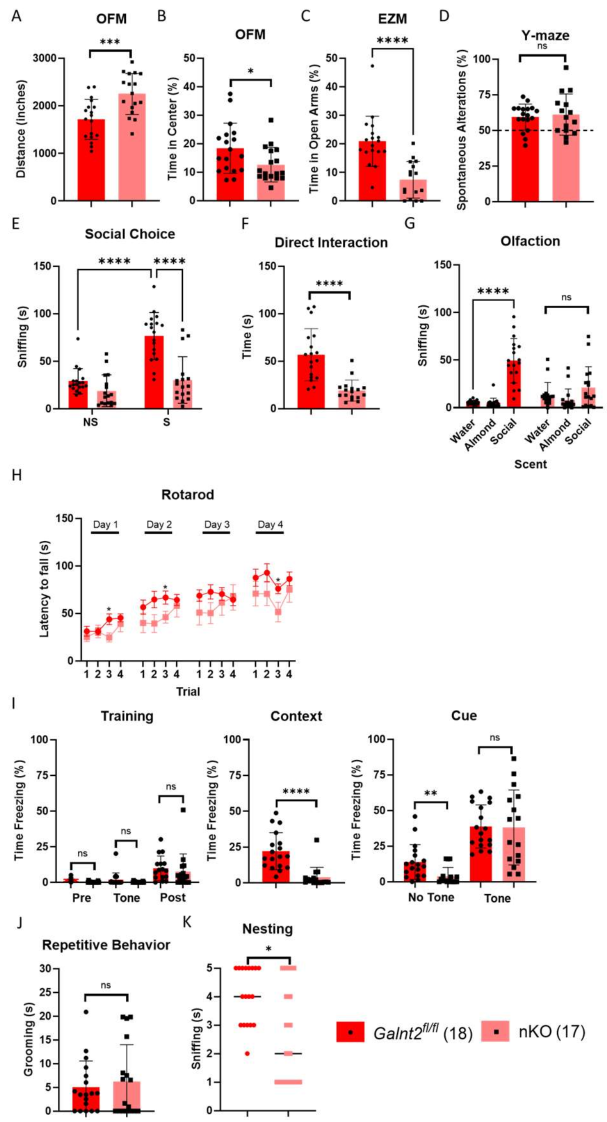


**Supplemental Figure 5. Female *Galnt2* nKO mice exhibit broad deficits across neurological motor, sensory, learning, and memory domains.** (A) Female nKO mice explore the OFM more than floxed littermates with (B) less time spent in the center of the OFM. (C) Female nKO spend less time in the open arms of the EZM. (D) Female nKO have intact short-term and spatial working memory in the Y-maze. (E) Female nKO mice spend less time than floxed littermates sniffing and (F) less time directly interacting with a novel stimulus mouse during the 3-chambered SC test, which could be due to (G) lack of social scent discrimination. (H) Female nKO fall more quickly from an accelerating, rotating rod than floxed littermates. (I) Female nKO mice spend significantly less time freezing compared with floxed littermates when returned to the fear-conditioning testing chamber (contextual) but (I) the same amount of time freezing upon hearing the footshock-associated tone (cue). (J) Female nKO mice spend no more time in repetitive grooming than floxed littermates. (K) Female nKO mice have trouble building well-formed nests. For all panels, floxed, $n$ = 18; nKO, $n$ = 17. Welch's unpaired, 2-tailed t-test, except as follows: 3-chambered SC assay and olfaction, two-way ANOVA with Fisher's Least Significant Differences test; rotarod, two-way matched values ANOVA with Fisher's Least Significant Differences test; nesting, unpaired Mann-Whitney test. *, $P < 0.05$, **, $P < 0.01$, ***, $P < 0.001$, ****, $P < 0.0001$. Data are represented as mean ± SD, except rotarod where errors bars represent ± SEM and nesting where the line indicates median.

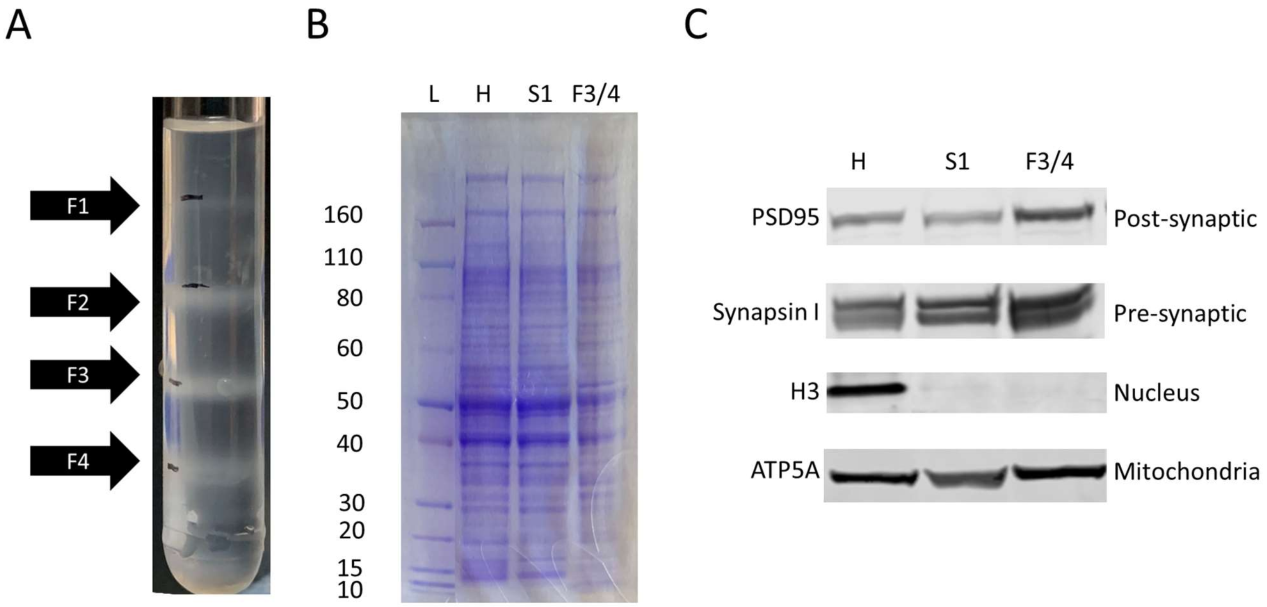


**Supplemental Figure 6. Percoll gradient isolation of synaptosomes.** (A) Representative picture of Percoll gradient following centrifugation and prior to fraction isolation. (B) Representative protein gel with Coomassie stain of aliquots from various stages of synaptosome isolation. (C) Representative western blots for PSD95, Synapsin I, H3, and ATP5A (post-synaptic density, pre-synaptic density, nucleus, and mitochondria, respectively) demonstrating enrichment of pre- and post-synaptic densities and depletion of nuclei in combined fractions 3 and 4. L, Ladder; H, cortical homogenate; S1, supernatant applied to the Percoll gradient; F3/4, Combined fractions 3 and 4 collected from the Percoll gradient.

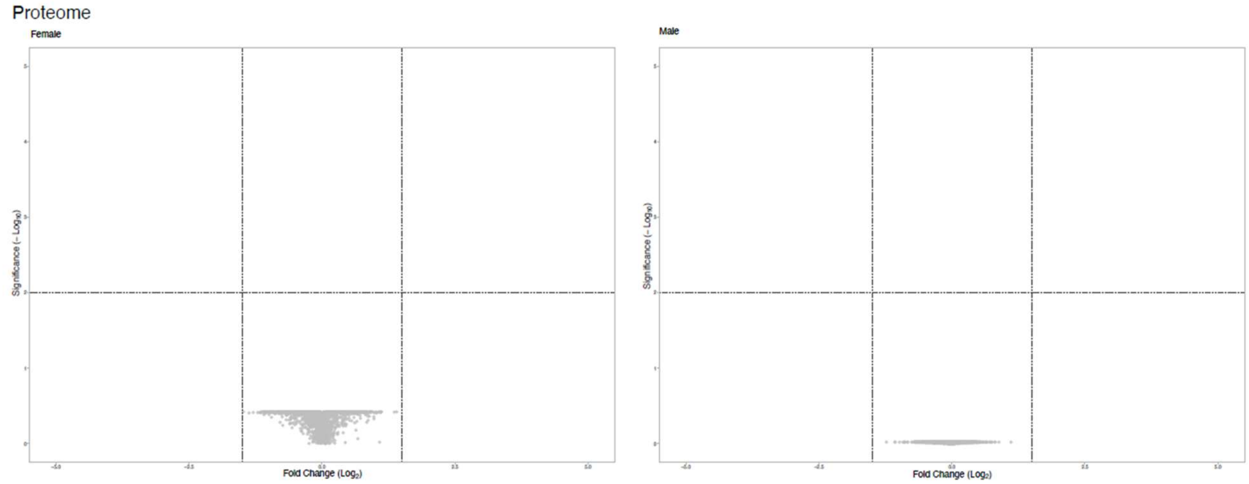


**Supplemental Figure 7. Contribution of GALNT2 to the cortical synaptosomal proteome.** Differential proteome between *Galnt2* nKO and floxed littermates is plotted as volcano plots of nKO/control fold change for female and male synaptosomes. Lack of pseudocoloring indicates no proteins with fold change > 3 and $P < 0.01$.

### SUPPLEMENTAL METHODS

#### Behavioral Assessments

*Open-field maze (OFM).* Locomotor activity was measured via an open-field maze test where mice were individually placed into, and allowed to explore, a 15” x 15” arena for a total of 30 min. A ceiling-mounted camera allowed for a video-tracking software (SmartScan 3.0) to real-time analyze the total distance traveled as well as the percent time spent in the center of the arena (defined as the central 25% of the total area). Activity was analyzed for the first 10 minutes of time in the maze.

*Elevated zeromaze (EZM).* The elevated zeromaze (San Diego Instruments; California, USA) consists of a circular shaped platform elevated 3 feet above the floor. Two opposite quadrants of the maze are enclosed (wall height, 12 inches), whereas the other two are open (wall height, 0.5 inches). Mice were placed in one of the closed quadrants and their movement traced over the course of 5 min. Analysis, including the quantification of percent of time spent in open arms and the number of entries, was performed manually using a stopwatch. An entry was defined as a transition from a closed to open arm, or vice versa, that involves all four paws. Mice that exited the maze prior to 5 minutes were excluded from analysis.

*Y-maze.* Spontaneous alternation behavior was measured on a Y-maze apparatus (San Diego Instruments; California, USA), composed of three arms (Arm A: 8in. x 5in. x 3in.; Arms B and C: 6in. x 5in. x 3in.). For testing, the mouse was placed in Arm C, facing the center, and allowed to freely explore the maze for 5 min. A spontaneous alternation was defined an entry into the arm less recently explored. Percent spontaneous alternation was calculated as the number of spontaneous alternations over the total number of entries. For example, the sequence C,B,A,B,C,B,A,C (starting in arm C) resulted in a percent spontaneous alternation of 4/6 = 67%. Mice with less than 15 total alternations in the 5 minutes were excluded from analysis.

*Three-chambered SC assay.* The SC test was carried out in a three-chambered apparatus that consisted of a center chamber and two end chambers. Before the start of the test and in a counter-balanced sequence, one end chamber was designated the social chamber, into which a stimulus mouse would be introduced, and the other end chamber was designed the nonsocial chamber. Two identical, clear Plexiglas cylinders with multiple holes to allow for air exchange were placed in each end chamber. A ceiling-mounted camera recorded the entire assay. In the habituation phase of the test (Phase I), the test mouse was placed in the center chamber and allowed to explore all three chambers for 10 min. In the social choice phase of the test (Phase II), an age-matched stimulus mouse (adult, gonadectomized A/J mice) was placed in the cylinder in the social chamber while a novel object was simultaneously placed into the other cylinder in the nonsocial chamber. During the subsequent 10 min social choice period, amount of time spent sniffing the social cylinder was documented by watching the video and recording with a stopwatch. In the direct social interaction test, the cylinders were removed simultaneously following the SC test, and the amount of time test and stimulus mice spent in direct contact (sniffing, allogrooming) was measured for a total of 5 min. If fighting persisted for more than several seconds, the mice were removed from the apparatus and excluded from analysis.

*Accelerating rotarod assay.* Mice were placed on an accelerating rotarod apparatus (Harvard Apparatus) for 16 trials (four trials a day for four consecutive days) with at least 15 min of rest between the trials. Each trial lasted for a maximum of 5 min, during which the rod accelerated linearly from 4 to 40 rpm. The amount of time for each mouse to fall from the rod was recorded for each trial.

*Context- and cue-dependent fear conditioning.* For the training day, mice were placed in individual chambers (Med Associates) for 2 min followed by a loud tone (85 dB, 2 kHz) lasting 30 s that co-terminated with a 2-s, 1.25-mA foot shock. Mice were left undisturbed for an additional 2 min 30 s in the chamber and then immediately placed back into their home cage. Freezing behavior, defined as no movement except for respiration, was determined before and after the tone-shock pairings and scored by FreezeScan NI version 2.00. To test for context-dependent learning, mice were placed back into the same testing boxes 24 hr later for a total of 5 min without any tone or shock, and again the total time spent freezing was measured. After 4 hrs, we tested for cue dependent fear memory by placing the mice into a novel chamber consisting of altered flooring, wall-panel inserts, and vanilla scent. After 2 min in the chamber, the cue tone (85 dB, 2 kHz) was played for a total of 3 min, and the total time spent freezing during the presentation of this cue tone was recorded.

*Olfaction.* Mice were tested for whether they could detect and differentiate odors in a modified habituation-dishabituation protocol (Yang, M. and Crawley, J.N. 2009). Mice were presented with cotton-tipped wooden applicators dipped in either water, vanilla, or swiped across the bottom of an unfamiliar opposite-sex social cage. Each stimulus was presented for 2 min with a 1-min inter-trial interval. Time spent sniffing was defined as when the animal was oriented with its nose 2 cm or closer toward the cotton tip. The assay was recorded with a horizonally-positioned camera and sniffing time manually recorded with a stopwatch.

*Repetitive behavior.* Mice were individually placed into a clean, home-cage like environment lined with bedding. After allowing 5 min for habituation, 10 min of activity was videotaped for each mouse. The duration of repetitive behavior, defined as grooming or digging, was scored manually using a stopwatch.

*Nesting.* Nest construction was evaluated using the metric as described previously (Deacon, R.M. 2006). 18 h after mice were singly housed with a cotton square nestlet (Ancare) and no other bedding material, nests were assessed for amount of nestlet material shredded, height, and shape and scored using the following metric: (1) nestlet not noticeably touched; (2) nestlet partially torn; (3) nestlet mostly shredded but with no identifiable nest site; (4) an identifiable but flat nest; and (5) a perfect nest with walls. A mouse whose cage flooded overnight was excluded from analysis.

**Molecular Studies**

*Protein gels.* Protein concentration was measured using a Bradford assay. Equal amounts of protein were loaded into each well of a 4%-12% Bis-Tris gradient gel (Invitrogen, 10 well, 1.0 mm) and run at 200 V for 50 minutes in MOPS buffer. Gel was Coomassie stained and destained following standard protocols.

*Western blots.* Proteins were transferred onto a PVDF membrane (0.45 µm pore size; Bio-Rad). The resulting membrane was blocked with a 1:1 solution of Odyssey blocking buffer (LICOR; 927-40100) and 1× PBS for 1 hour at room temperature. Primary antibodies used were rabbit anti-PSD95 (Cell Signaling, 3450, 1:1000), rabbit anti-Synapsin I (Millipore, AB1543P, 1:1000), rabbit anti–Histone H3 (Millipore, 06-755, 1:1000), and mouse anti-ATP5A (Abcam, ab14748, 1:1000). Secondary antibodies (LI-COR) used were goat anti-rabbit IgG (IRDye 800CW, catalog 926-32211, 1:10000) and goat anti-mouse IgG (IRDye 680LT, catalog 926-68020, 1:10000), incubated for 60 minutes at room temperature. Standard protocols were used for the Odyssey Infrared Imaging System (LI-COR) for protein visualization and quantification.
